# Developmental dynamics of sex reprogramming by high incubation temperatures in a dragon lizard

**DOI:** 10.1101/2021.09.23.461594

**Authors:** Sarah L. Whiteley, Clare E. Holleley, Arthur Georges

## Abstract

In some vertebrate species, gene-environment interactions can determine sex, driving bipotential gonads to differentiate into either ovaries or testes. In the central bearded dragon (*Pogona vitticeps*), the genetic influence of sex chromosomes (ZZ/ZW) can be overridden by high incubation temperatures, causing ZZ male to female sex reversal. Previous research showed ovotestes, a rare gonadal phenotype with traits of both sexes, develop during sex reversal, leading to the hypothesis that sex reversal relies on high temperature feminisation to outcompete the male genetic cue. To test this, we conducted temperature switching experiments at key developmental stages, and analysed the effect on gonadal phenotypes using histology and transcriptomics. We found sexual fate is more strongly influenced by the ZZ genotype than temperature. Any exposure to low temperatures (28°C) caused testes differentiation, whereas sex reversal required longer exposure to high temperatures. We revealed ovotestes exist along a spectrum of female-ness to male-ness at the transcriptional level. We found inter-individual variation in gene expression changes following temperature switches, suggesting both genetic sensitivity to, and the timing and duration of the temperature cue influences sex reversal. These findings bring new insights to the mechanisms underlying sex reversal, improving our understanding of thermosensitive sex systems in vertebrates.

## Introduction

Sex determination in vertebrates exists on a continuum spanning from genetic sex determination (GSD) to temperature dependent sex determination (TSD) (1). Some species possess sex chromosomes with a thermal override that can cause sex reversal. In the case of the Australian central bearded dragon *Pogona vitticeps*, the most well studied reptile with sex reversal, genetic males (ZZ sex chromosomes) incubated at high temperatures (>32°C) undergo sex reversal so the animal develops as a female despite being genetically male (2,3). As these sex reversed females (ZZf) are reproductively viable, mating of ZZf and ZZm individuals yields offspring whose sex is determined solely by temperature in the absence of the W chromosome (2,3). Despite this being akin to that observed in TSD species, where sex is determined in the absence of sex chromosomes, sex reversal in *P. vitticeps* differs in several important ways. The offspring of sex reversed mothers inherit ZZ chromosomes, and reversal of their offspring may be more sensitive to temperature than ZZ offspring of ZW mothers (2). Some individuals do not sex reverse at high temperatures, suggesting interindividual propensity for sex reversal. Variability in rates of sex reversal also exists at the population level (4). Both thermally-induced sex reversal and TSD require influential cells of the bipotential gonad to sense and transduce temperature to epigenetic changes that ultimately govern sex determination and differentiation (5).

Understanding the mechanisms of TSD has remained elusive despite decades of attention following the discovery of TSD in the African dragon *Agama agama* (6). The most recent proposition for how an environmental cue can cause cellular changes that ultimately determine sexual fate invokes calcium and redox signalling at the head of the regulatory cascade (5). This CaRe model predicts involvement of genes that govern the capacity of the cell to sense and respond to environmental changes and to subsequently modulate expression of genes in ubiquitous signalling pathways. The modulation of these signalling pathways leads to epigenetic processes (e.g. action of chromatin modifier genes) that influence the expression of sex genes (7). CaRe mechanisms have been implicated in the TSD cascades of several species (5), and most recently in the gene expression patterns associated with sex reversal in embryonic *P. vitticeps* (8). Four genes are consistently associated with sex reversal and TSD systems namely, *JARID2, KDM6B, CIRBP* and *CLK4* (7–12). Chromatin remodelling genes *JARID2* and *KDM6B* regulate gene expression through modulation of methylation marks on lysine 27 on histone 3 (H3K27) (13,14). In *Trachemys scripta KDM6B* demethylation of the *DMRT1* promoter is necessary for male development (7). *CIRBP* is an environmentally sensitive gene that is activated by temperature, and splicing of this gene has been shown to be controlled by *CLK4*, which is itself temperature sensitive (11,12). *CLK4* has also been shown to regulate the splicing of *JARID2* in two TSD turtles (12).

In TSD species, the epigenetic mechanisms responsible for sensing temperature and affecting gene expression can only influence sex during a window of embryonic development within which gonadal fate is responsive to temperature, known as the thermosensitive period (TSP). After the TSP, gonadal fate has been irreversibly determined (15). The boundaries of the TSP are typically identified using temperature switching experiments, whereby eggs are switched between male and female producing temperatures at different embryonic stages during development (15). A less common approach is to apply hormone inhibitors at different developmental stages to determine when the embryo becomes insensitive to its affects (16). However, the timing of the TSP is not necessarily precise. For example, in *Alligator mississippiensis* the TSP occurs from embryonic stages 21 to 24 (ca 30-45 d post lay at 29-31°C) (17). Subsequent experiments showed that much earlier incubation conditions experienced around stage 15 can also influence sex (18). Incubation at temperatures intermediate to the male and female producing temperatures (at the pivotal temperature) typically yield a mixture of the two sexes, presumably because temperature is equivocal in its influence allowing subtle genetic predisposition to prevail (19). In TSD systems the presence of such underlying genetic predisposition is obscured by the dominant effect of temperature at all but a very narrow thermal range. In systems that display sex reversal, as in *P. vitticeps*, the complexity of gene-environment interactions involving temperature is further compounded by the presence of sex chromosomes (20).

The development of ovotestes is one effect of temperature on gonad differentiation. Ovotestes are a gonadal phenotype with both male and female traits (21). In TSD species, ovotestes are rarely observed in embryos, and when they occur are likely a result of hormone balances at intermediate incubation temperatures (22,23). In contrast, temperature induced sex reversal in *P. vitticeps* is observed as the temporary presence of ovotestes at stage 9 in approximately 40% of reversing embryos (21). Ovotestes in ZZ/ZW species are thought to be caused during sex reversal by the competing influences of male sex chromosomes and feminizing incubation temperatures (21). As such, sex reversal is hypothesised to be a process by which the feminizing influence of temperature must override that of the masculinizing influence of the sex chromosomes (21).

To test the hypothesis that sex reversal involves over-riding the male sex determining signal, we switched incubation temperatures between 28°C where sexual phenotype and genotype are concordant and 36°C where sex reversal of ZZ individuals to a female gonadal phenotype is predominant. Switches were conducted at three developmental stages (Fig. 1) ranging from bipotential to differentiated (21,24), and the gonadal phenotypes were analysed with histology and RNA-sequencing. With both morphological and gene expression data, the complex effects of temperature switching on sex determination and differentiation in *P. vitticeps* were revealed. We showed the timing and duration of the feminising high temperature cue that was required to override the male signal and initiate sex reversal. We profiled the gene expression characteristics of ovotestes, and temperature response genes, many of which have been implicated in the CaRe model (5) or in TSD (7,11,12). This research provides new insights to the mechanisms of sex reversal in *P. vitticeps*, but also to vertebrate thermolabile sex determination more broadly.

**Figure 1:**
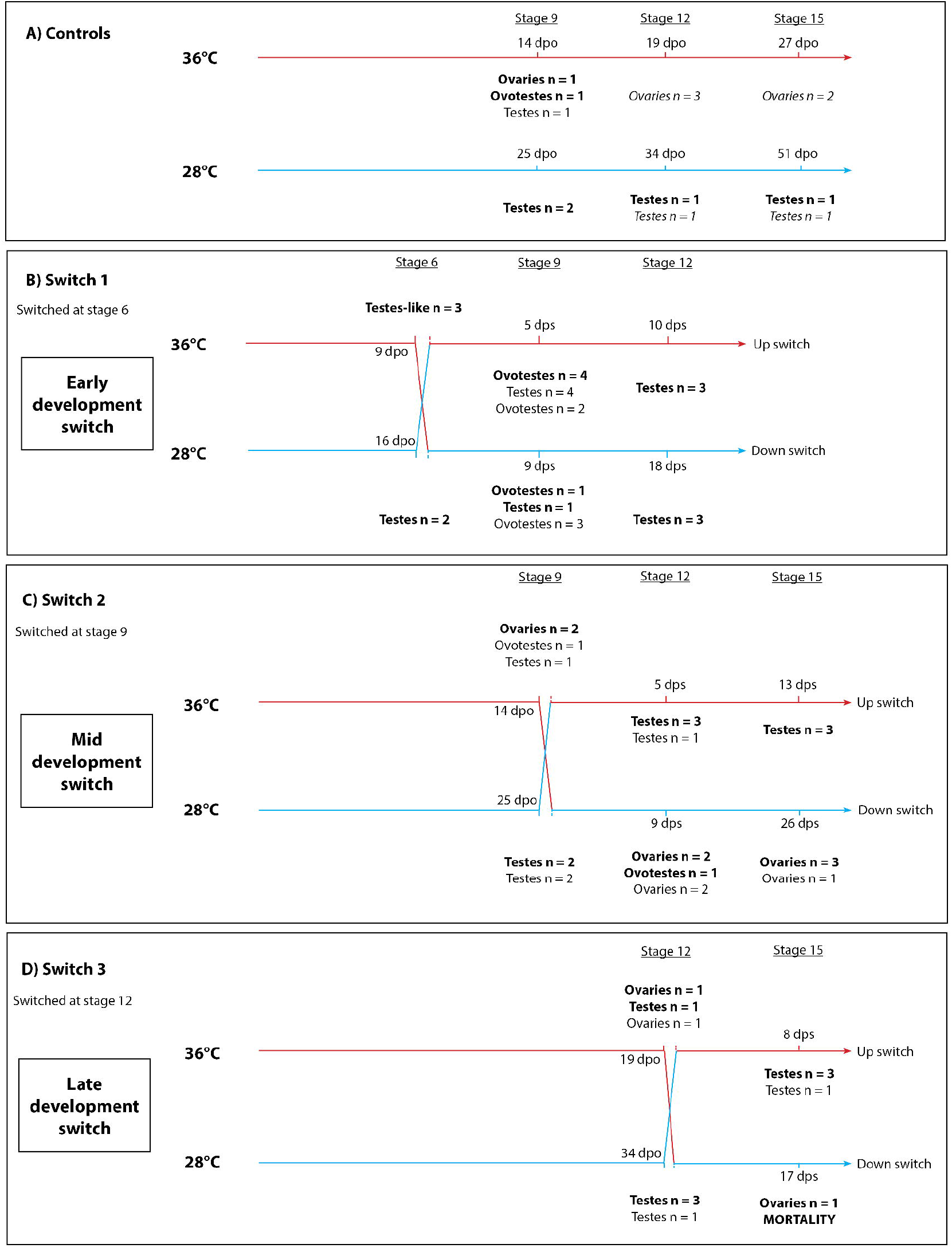
Experimental design showing the control treatment (A) and the three temperature switch treatments (B-D). Gonadal phenotypes were determined using RNA-seq and histology (bold), histology only (normal text) and for a subset of the control samples, RNA-seq only (italic text).Abbreviations: dpo, days post oviposition; dps, days post switch. A mortality rate of 80% occurred in the late development switch following a down switch at stage 12 to 36°C to 28°.

## Results

### Temperature switching and gonadal phenotype

#### Control Experiments

Incubation at 28°C without a switch resulted all individuals developing testis at all developmental stages, as would be expected from ZZ individuals (Fig. 1A, D) (2). Incubation at 36°C yielded gonads with testis-like gonads at Stage 6, ovotestes and ovaries at stage 9, and ovaries at stages 12 and 15 (Fig. 1A, D). Testis-like gonads were characterised by rudimentary seminiferous tubules akin to those observed in ovotestes, but without a thickened cortex (Fig. 2). These results suggest that testes are the default developmental outcome for ZZ individuals, influenced by high temperature to become ovaries.

**Figure 2:**
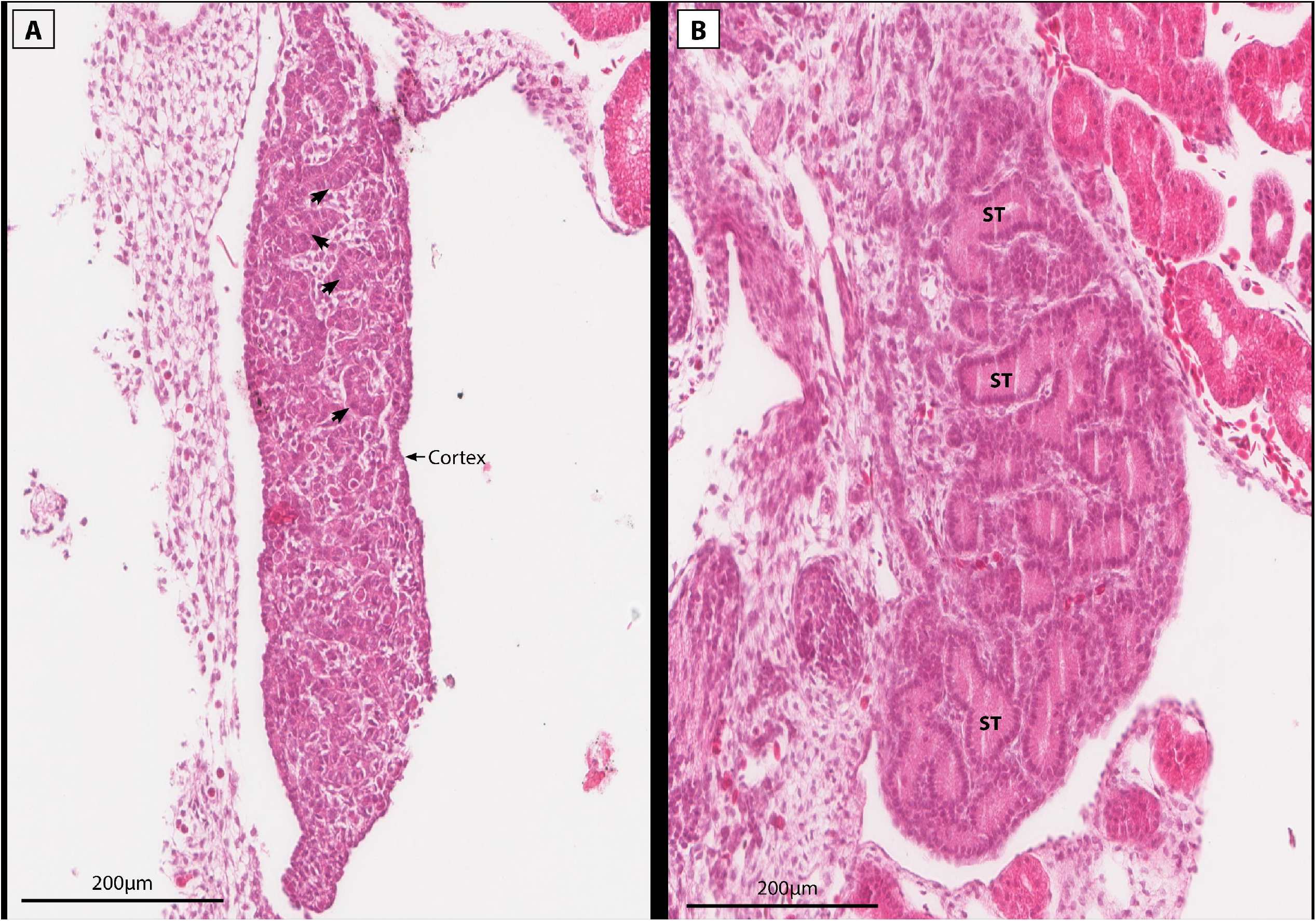
Sections of two stage 6 embryonic gonads from the same clutch with a testes-like phenotype (A) and normal testes (B). The testes-like phenotype was produced at 36°C, and is characterised by having an elongated shape typical of a bipotential gonad, with rudimentary seminiferous tubules typical of ovotestes (arrows), but having only a thin cortex layer. For comparison, a typical testes is displayed showing no cortex layer and well developed seminiferous tubules (ST).

#### Switch 1 (Stage 6)

The gonadal condition at stage 6, immediately before the switches from 28°C to 36°C (hereafter up-switched) and 36°C to 28°C (hereafter down-switched), was described under the control experiments. Embryos up-switched at stage 6 had ovotestes or testes by stage 9, and testes by stage 12. Thus, exposure to 28°C for the period leading up to stage 6 was sufficient to irreversibly determine a male sexual fate, one that could not be reprogrammed by subsequent exposure to the sex reversing temperature of 36°C.

Embryos down-switched at stage 6 had ovotestes or testes by stage 9 and testes by stage 12 (Fig. 1B). This suggests that exposure to sex reversing temperature of 36°C in the period leading up to stage 6 was not sufficient to irreversibly overcome the reprogramming of male sexual fate by subsequent exposure to 28°C.

#### Switch 2 (Stage 9)

Embryos up-switched at stage 9 had ovotestes or testes by stage 12, maintained as testes to stage 15. Thus, exposure to 28°C for the period leading up to stage 9 was, as expected from the results of Switch 1 experiments, sufficient to determine irreversible male sexual fate.

Embryos down-switched at stage 9 had ovotestes or ovaries by stage 12 and ovaries by stage 15 (Fig. 1B). This suggests that exposure to sex reversing temperature of 36°C in the period leading up to stage 9 was sufficient to irreversibly determine female fate, and overcome the programming of male sexual fate despite subsequent exposure to 28°C.

#### Switch 3 (Stage 12)

In the context of the results of the above experiments, the results of the third switch experiment were expected (Fig. 1C). The embryos up-switched at stage 12 had testes by stage 15, and those down-switched at stage 12 had ovaries by stage 15, though mortality was high in this latter experiment.

The broad interpretation of these results is that the default developmental program of sexual fate in ZZ individuals is male, as one would expect, and that sexual fate is more strongly influenced by the ZZ genotype than temperature. Indeed, early exposure to 28°C in any of our experiments led to a male gonadal phenotype. Reprogramming of sexual fate under the influence of temperature was not observed until eggs were incubated at 36°C to stage 9. The thermosensitive period for sex reversal thus begins after stage 6 and at or before stage 9.

### Differential gene expression analysis

These data provide unique opportunities to compare combinations of developmental stage, temperature, and phenotype that are not normally possible. Principal component analysis (PCA) on normalised read counts detected an outliner sample (Fig. 3A) that was removed from further analysis. As expected based on the morphological characteristics of ovotestes, the samples with ovotestes were distributed between ovaries and testes along the PC2 (Fig. 3B).

**Figure 3:**
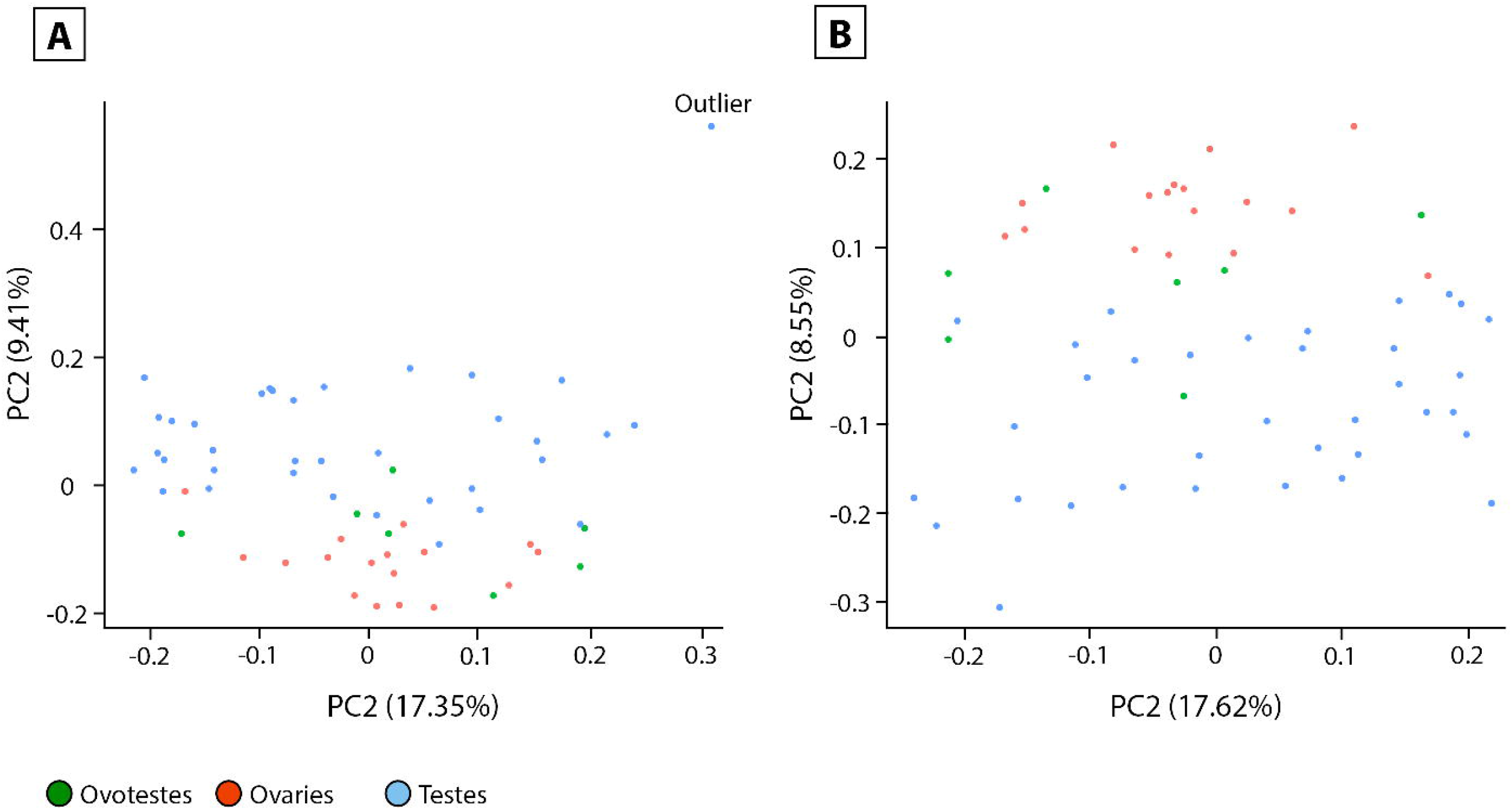
PCA plots for the complete dataset coloured by phenotype (A) and with an outlier sample (sample ID 3344zz_18_1_19) removed in panel B. The principle component analysis was conduced on normalised read counts.

#### Testis-like gonads vs testes (Stage 6)

In the switch 1 regime, differentially expressed genes between the testes-like gonads observed in 36°C embryos at stage 6 (control and switch experiments) compared with their 28°C incubated counterparts with normal testes, revealed genes that are associated with temperature at the same developmental stage, as well as the genes upregulated in the testes-like phenotype observed at 36°C (Supplemental Table S1).

A variety of genes with sex-related functions are differentially expressed between the testes-like gonads and the normal testes. In the 28°C testes, a newly discovered testes-specific gene, *TEX33*, which is involved in spermatogenesis was upregulated (25). *GCA* (grancalin) was also upregulated. Grancalin has previously been implicated in GSD female development in *P. vitticeps*, but the data here suggests a more generic role in reproduction that is not sex specific (8). The testes-like gonads displayed upregulation of a mix of both male and female related genes compared to the testes. These included male specific genes *SLC26A8* (which was the most differentially expressed), *SRD5A1*, and *SOX9*. In mammals, *SLC26A8* is required for proper sperm functioning (26). *SRD5A1* catalyses testosterone into the more potent dihydrotestosterone, and *SOX9* is a canonical testes gene. At the same time, *WNT9a* was also upregulated, a gene in the *Wnt* signalling pathway associated with ovarian development. Also upregulated was *PIAS1*, a gene that can downregulate *SOX9* in mammals (27).

Various environmentally responsive genes were upregulated in the 36°C testes-like gonads. *CIRBP*, a gene associated with sex reversal in *P. vitticeps* and other TSD species, was upregulated. A member of the Jumonji family, *JMJD6*, which demethylates H3K4 and H3K36, was upregulated, as was *KDM4b*, which demethylates H3K9. Mitogen activated protein kinases, *MAP3K13, MAPKAPK5, MAPK1*, which mediate cell signalling pathways in response to environmental stress, were upregulated (28). Other genes related to calcium and redox responses (CaRe) were also differentially expressed between the two groups. These included *MICU, GLRX2, TXNDC5, OXSR1, C2CD2*, and *PIDD1* (29).

#### Testes at 28°C vs testes at 36°C (Stage 12)

The early developmental switch regime produced testes at both 28°C and 36°C at stage 12. A suite of genes were differentially expressed between these two groups (Supplemental Table S2). In the 28°C testes, *POMC* was upregulated. This is an interesting result because in *P. vitticeps, POMC* upregulation has been previously associated with adult sex reversed females (10). *CATSPER4* was highly upregulated in 28°C testes, and is associated with essential sperm functions. *RARG* a retinoic acid receptor that is normally associated with ovarian development (30) was also upregulated at 28°C, as was *GCA*, a gene previously associated with ZW female development in *P. vitticeps* (8). Also upregulated were spliceosome genes *SNU13* and *PRPF38A* and DNA repair genes *SUMO2* and *UBE2N*. A suite of environmental stress genes was upregulated at 36°C. These included heat shock genes *DNAJB14* and *DNAJC13*, NF-kB and STAT pathway genes *RelB, IKBKE, and STAT6*, mitogen activated protein kinase *MAP3K13* and antioxidant regulator *TXNDC11*. Well known chromatin remodelling genes *KDM6B* and *JARID2* were upregulated, as was *KATA2*, a lysine acetyltransferase. Calcium regulating genes T*RPV1, CACNB3, CADPS2* and *CABP1* were also upregulated in 36°C testes.

#### Testes at 28°C vs ovaries at 36°C (Stage 9)

In the mid development switch regime, at stage 9 ovaries (36°C) and testes (28°C) display sex specific expression (Supplementary Table S3). In ovaries, aromatase, *CYP17A1, GATA2*, and *FOXL2* were upregulated, while in testes *SOX9, WIF1, HSD17B3, CYP26B1* were upregulated. However, genes normally associated with the opposite sex were also upregulated. At 28°C these included *FZD3, FRZB*, and *WNT6* which are associated with the *Wnt* and *Beta catenin* pathways (31,32). In ovaries at 36°C, *SRD5A2*, a gene that converts testosterone to dihydrotestosterone(33), was upregulated. In the 36°C ovaries various CaRe related genes were upregulated, such as *CIRBP, TRPV1, JARID2, KDM6B, CACNA1I* and *CACNA1C, RALY, OSGIN1, DDIT4* and *GADD45A*. These results suggest that the underlying transcriptional profile can exhibit expression of genes associated with both sexes regardless of the phenotype. There are two possible explanations for this result; either some flexibility remains at the transcriptional level despite sex specific gonadal phenotypes, or gene expression may be able to change more rapidly than the cellular structures in the gonad.

### Expression trends of temperature associated genes

The chromatin remodelling genes *JARID2* and *KDM6B*, and the temperature sensitive *CIRBP*, were highly expressed during sex reversal, and likely play an important role in initiating the female pathway (8). Detailed analysis of the expression of these three genes showed they all exhibited similar expression patterns. *JARID2, KDM6B* and *CIRBP* all had higher expression at 36°C regardless of prior switch history, and expression decreased following up-switching. This suggests that these three genes are capable of rapid responses to environmental change. The opposite pattern occurs following up-switch, which caused an increase in expression. In all three genes there was considerable variation in expression levels between different individuals. Expression of these genes was significantly higher at 36°C compared to 28°C, but expression levels did not differ significantly between the three different phenotypes (ovaries, testes, and ovotestes).

*KDM6B* (Fig. 4) exhibited the highest expression level in the switch 1 regime after the up-switching, where two ovotestes samples exhibited high expression levels at stage 9 following up-switching at stage 6. Interestingly, another two ovotestes from the same regime had significantly lower expression, suggesting inter-individual variation in *KDM6B* expression changes in response to temperature.

**Figure 4:**
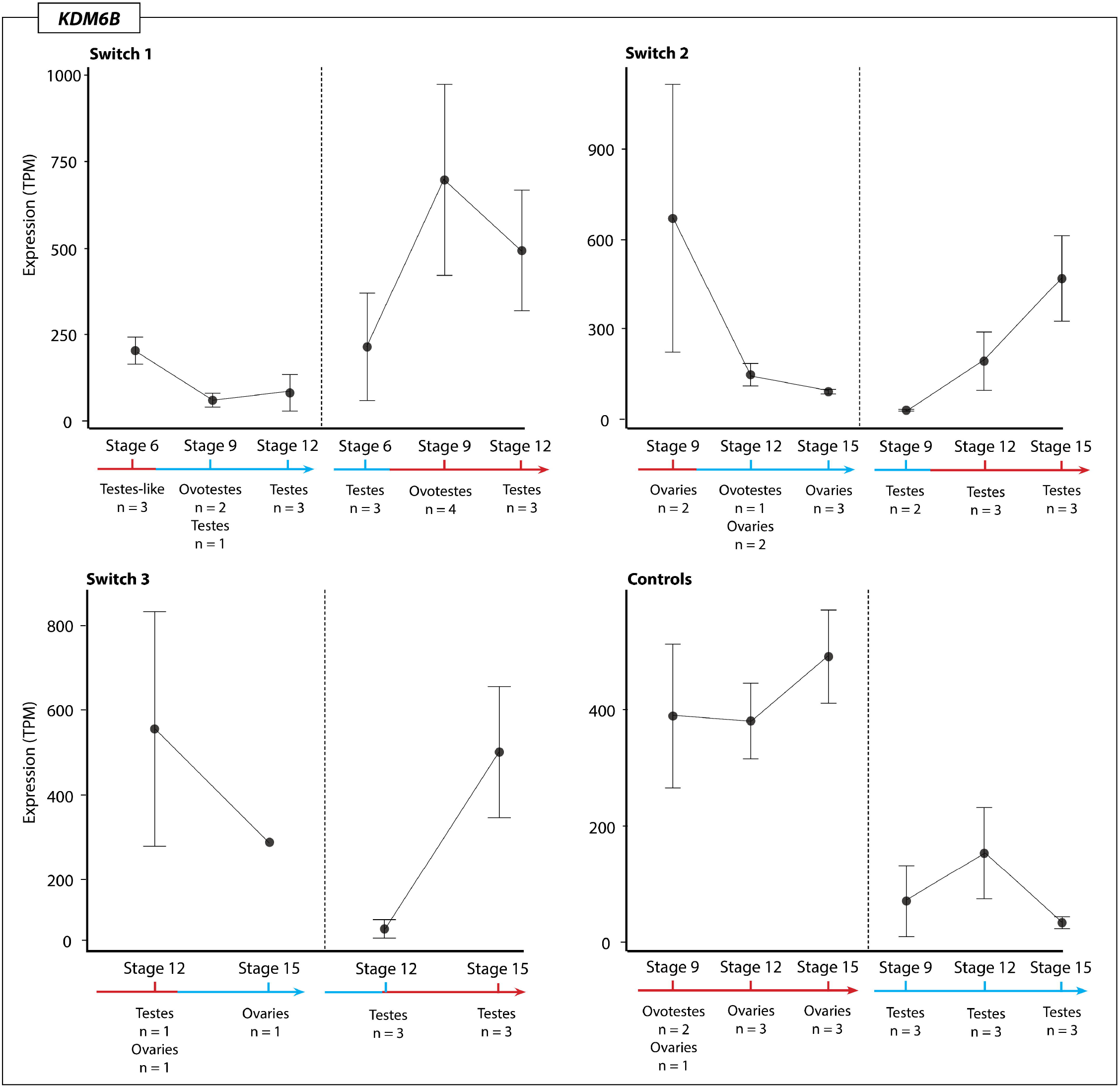
Expression (TPM, transcripts per million) of KDM6B for each temperature switch regime and controls groups at each sampling point regardless of phenotype. The variation in expression levels, mostly driven by the presence of different phenotypes, is reflected in the standard error bars.

*KDM6B* expression in switches between 28°C to 36°C for the three phenotypes (ovaries, testes, and ovotestes) was significantly higher at 36°C, but did not significantly differ between the phenotypes that were observed across the whole dataset. Taken together, these results suggests that *KDM6B* expression is sensitive to temperature and increases expression in response to higher incubation temperatures. *JARID2* (Fig. 5) exhibits similar patterns to *KDM6B*, though overall is more lowly expressed. Down-switching caused a decrease in *JARID2* expression, while up-switching caused increased expression. Compared to both *JARID2* and *KDM6B, CIRBP* was very highly expressed under all temperature conditions, and showed the same tendency to increase following exposure to 36°C and decrease following exposure to 28°C (Fig. 6).

**Figure 5:**
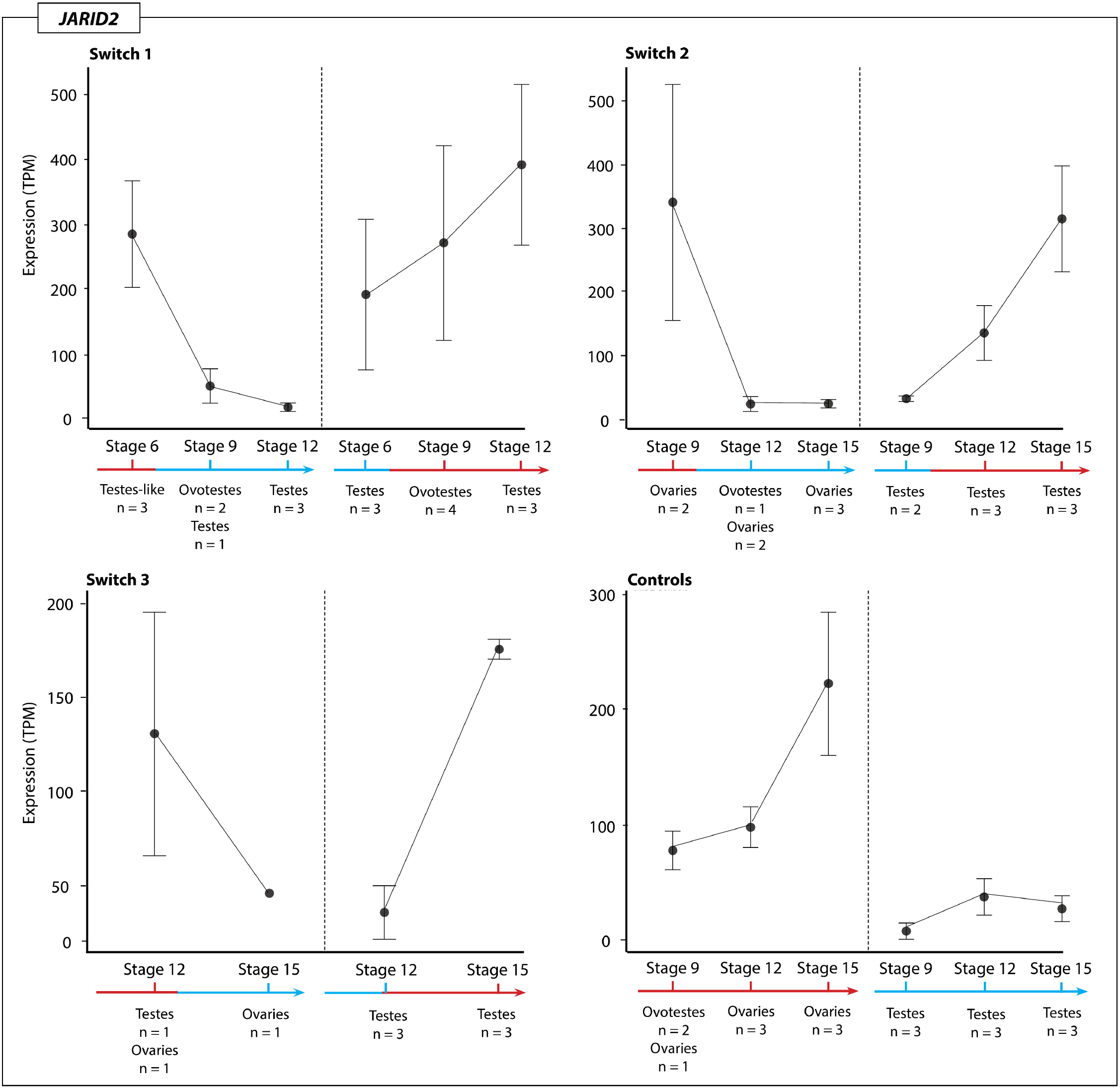
Expression (TPM, transcripts per million) of JARID2 for each temperature switch regime and controls groups at each sampling point regardless of phenotype. The variation in expression levels, mostly driven by the presence of different phenotypes, is reflected in the standard error bars.

**Figure 6:**
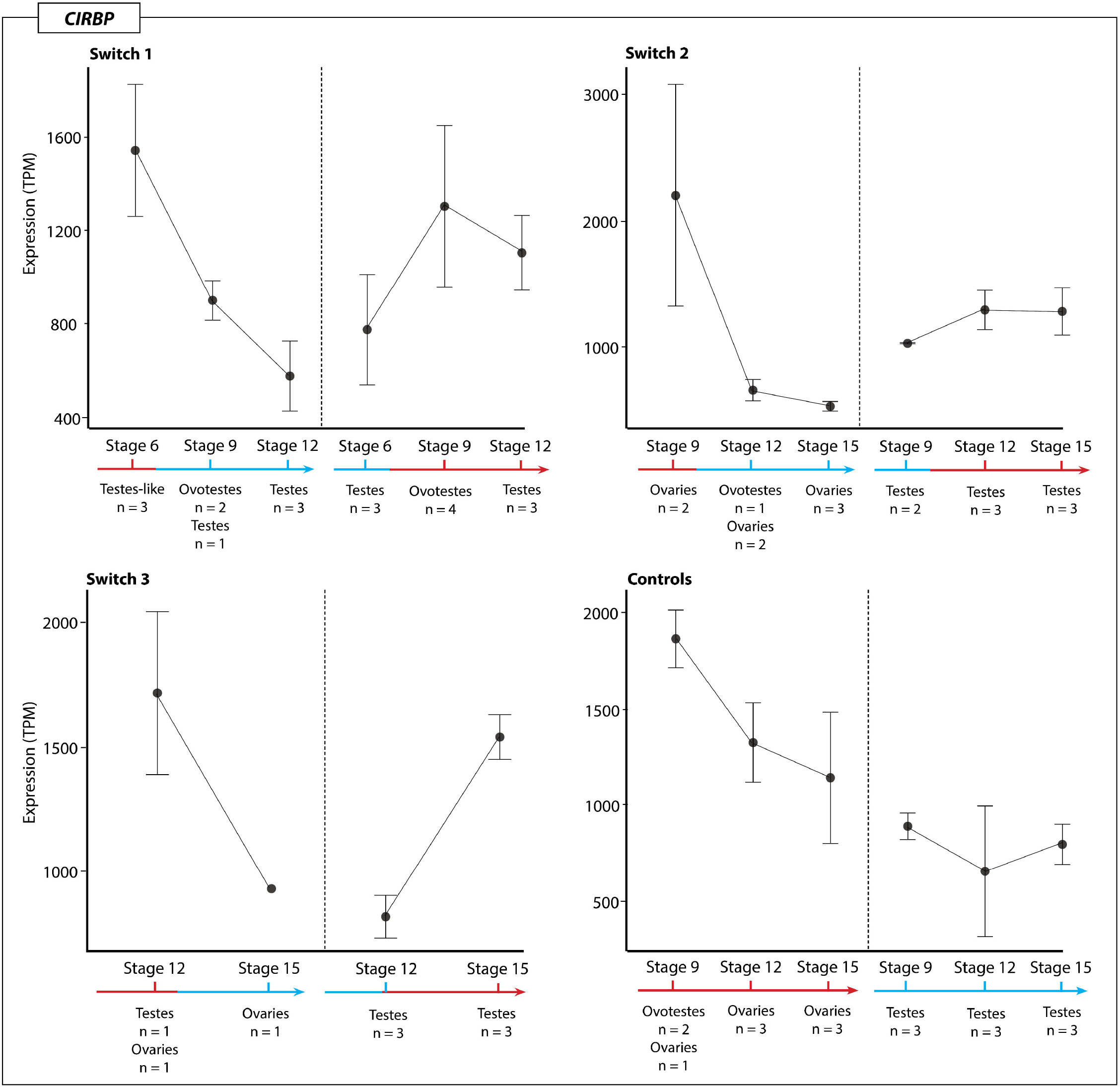
Expression (TPM, transcripts per million) of CIRBP for each temperature switch regime and controls groups at each sampling point regardless of phenotype. The variation in expression levels, mostly driven by the presence of different phenotypes, is reflected in the standard error bars.

### Transcriptional Profile of Ovotestes

As expected based on previous research (21) conducted at constant incubation temperatures, ovotestes occurred at only at stage 9 in the 36°C control constant incubation temperature experiments. Temperature switching experiments did affect the frequency of observing ovotestes, and also affected when during development they were observed. Specifically, ovotestes were observed in stage 9 embryos immediately following early developmental switches, regardless of the direction of the change (up-switch n = 6, ovotestes, down-switch n= ovotestes; Fig. 1B). Ovotestes were also observed in one stage 12 embryo individual subjected to a mid-development down-switch at stage 9 (Fig. 1C).

#### Ovotestes at 36°C vs ovotestes at 28°C (Stage 9)

One hundred genes were upregulated in stage 9 ovotestes generated after down-switching at stage 6 compared with those produced after up-switching. Genes upregulated following down-switching included antioxidant gene *TXNDC5, NOVA1* and *PRPF31* associated with splicing regulation in mammalian systems, and *CTBP2* associated with sex reversal in humans. *GCA*, a gene previously associated with *Pogona* ZWf female development at 28°C (8), was also upregulated (Supplemental Table S4).

Fifty two genes were upregulated in ovotestes generated after up-switching at stage 6. They included heat shock genes *DNAJB14* and *HSP90B1*, suggesting that switching to a high incubation temperature initiates the heat shock response. Also upregulated was *PDCD7*, a gene associated with splicing regulation and apoptosis, *RSF1* associated with chromatin remodelling and DNA repair, and a calcium signalling gene *CAMK2N1* (Supplemental Table S4).

#### Control ovotestes vs stage 6 up-switched ovotestes (Stage 9)

The transcriptional profiles of ovotestes examined at stage 9 following up-switching at stage 6 were compared to those of ovotestes at stage 9 in the control experiments. Genes upregulated in the up-switched ovotestes included *CAMK2N1* and *ATP2A1* both associated with calcium regulation, *PAX8* associated with thyroid hormone signalling, *NOX4* which neutralises reactive oxygen species and *PRKCZ* which regulates stress pathways in response to environmental stimuli. In the control ovotestes, the following were upregulated: (a) mitogen activated protein kinases *MAPKAP1* and *MAPK6*, (b) various circadian related genes including *RelB, CIPC* and *PER1*, (c) sex related genes *FZD5* (*Wnt* signalling), *CYP17A1* and *SHBG*, (d) inhibitor of the NF-κB pathway, *NFKBIE* and (e) *GADD45A*, an environmental stress response gene (Supplemental Table S5).

#### Stage 9 ovotestes vs stage 12 testes (36°C)

Comparison of ovotestes at stage 9 post-up-switch at stage 6 with the testes sampled later at stage 12 (switched at stage 6) can yield insights into the genes involved in resolving the gonad into testes rather than ovaries at the same incubation temperature (Supplemental Table S6). Genes upregulated in stage 9 ovotestes included stress response genes *CRH, CHR2, STK24, GADD45A, GSTP1*, and *MSRB1*. Thermosensitive calcium channel *TRPM5* was upregulated, as was calcium signalling gene *CAMK2N1*. Hormone related genes *CYP17A1, CYP19A1, DHRS3*, and *DHRS12* were upregulated. In the stage 12 testes produced after the temperature switch at stage 6, upregulated genes included various sex related genes including *HSD17B3, SHBG, SOX14, PAX7, FRZB*, and *WNT6*. Stress related genes were also upregulated including *DHRS7C, DNAJC11, HSPB2, PIDD1, DYRK3* and *TXNDC11*. Various calcium channels and sensors were upregulated including *CACNG1, CACNA1A, CCBE1*, and *CASQ1*

#### Stage 9 ovotestes vs stage 9 ovaries (36°C)

Comparison of ovotestes at stage 9 post up-switch at stage 6 (early developmental switch regime) with ovaries at stage 9 at 36°C (mid developmental switch regime) can reveal the differences between the phenotypes at the same stage and temperature, including the influence of the temperature switch (Supplemental Table S7). Genes upregulated in the pre-switch stage 9 ovaries included sex related genes *CYP17A1, SRD5A2, FZD1*, and *TDRP*, and stress related genes *ERO1B, OSGIN2, RRM2B*. Genes upregulated in ovotestes produced following up-switch, included *GCA*, circadian clock gene *CIART*, mitogen activated protein kinases *MAPK13* and *MAP3K9*, and thermosensitive calcium channel *TRPM3*.

#### Sex specific gene expression in ovotestes

When assessing the expression levels of sex specific genes (male associated genes *SOX9, AMH, NR5A1*, and female associated genes *CYP19A1, CYP17A1*, and *Beta-catenin)*, considerable variation across individuals was apparent (Fig. 7). For some genes, expression levels between individuals from the same switch regime, and even from within the same clutch, could be orders of magnitude different. For example, from the samples in the switch 1 regime, Sample B (sample ID 3603zz_18_1_7) expressed *SOX9* and *AMH* well above the mean, and well above that of its counterparts from the same clutch (Fig. 7A). Sample B also had expression well above the mean for *SOX9*, but not for *NR5A1*. For *CYP19A1* and *CYP17A1*, expression was nearly 0, but then expression for *Beta-catenin* was above average (Fig. 7B). This shows that an individual can radically differ in expression levels of genes compared to other individuals from the same clutch, including in expression levels of sex related genes.

**Figure 7:**
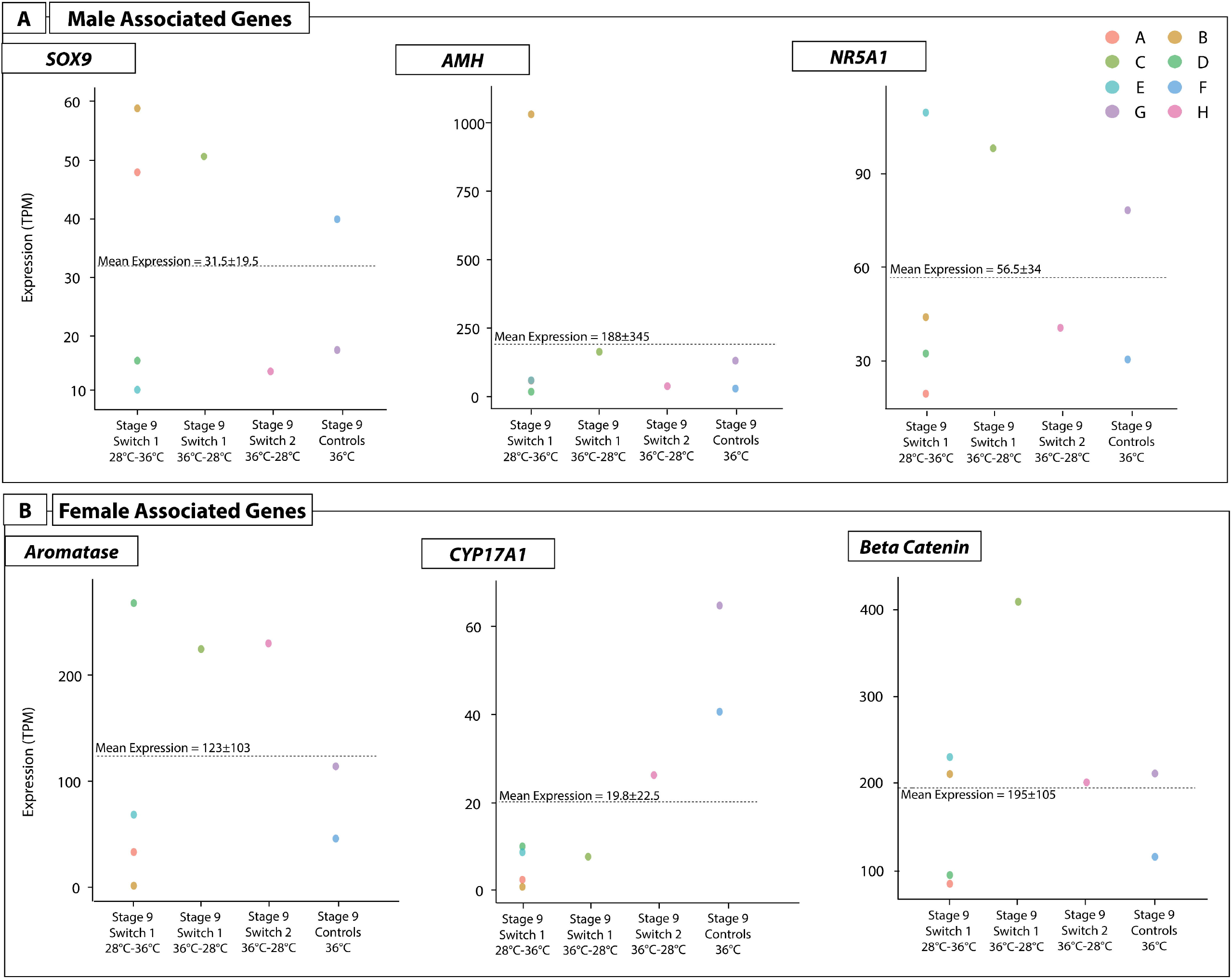
Expression (TPM, transcripts per million) of three male and three female sex-associated genes for all samples with ovotestes produced from different switching conditions (x-axis). Each point represents an individual (by colour) to depict the range of expression values different individuals can exhibit for a given gene. Sample IDs correspond to those provided in Supplementary File S13, and are as follows: A) 3603zz_18_1_4, B) 3603zz_18_1_7, C) 3603zz_18_1_8, D) 3603zz_18_2_4, E) 3632zz_18_2_9, F) 3632zz_18_2_19, G) 3632zz_18_2_20, H) 3232zz_18_1_19.

The ovotestes phenotype appears to exist along a spectrum, where at one end they share more similarities with ovaries than testes, at the other they are more similar to testes than ovaries, and in the mid-range they are a true intermediate between ovary and testis. This spectrum was evident in gene expression levels. It was also evident on histological examination in this study and previous experiments (21).

#### Temperature associated gene expression in ovotestes

Examination of the expression levels of genes associated with temperature response (*KDM6B, JARID2, CIRBP* and *CLK4*) reveal similar inter-individual variation occurred as in the expression of sex associated genes (Fig. 8). *CIRBP* exhibited the highest expression levels, and the greatest variation in expression. For example, in the stage 9 samples up-switched at stage 6, expression of *CIRBP* was several orders of magnitude different between individuals, ranging from 500 to over 2000 transcripts per million 9 (Fig. 8). Sample E was the only individual to have above average expression for these four genes.

**Figure 8:**
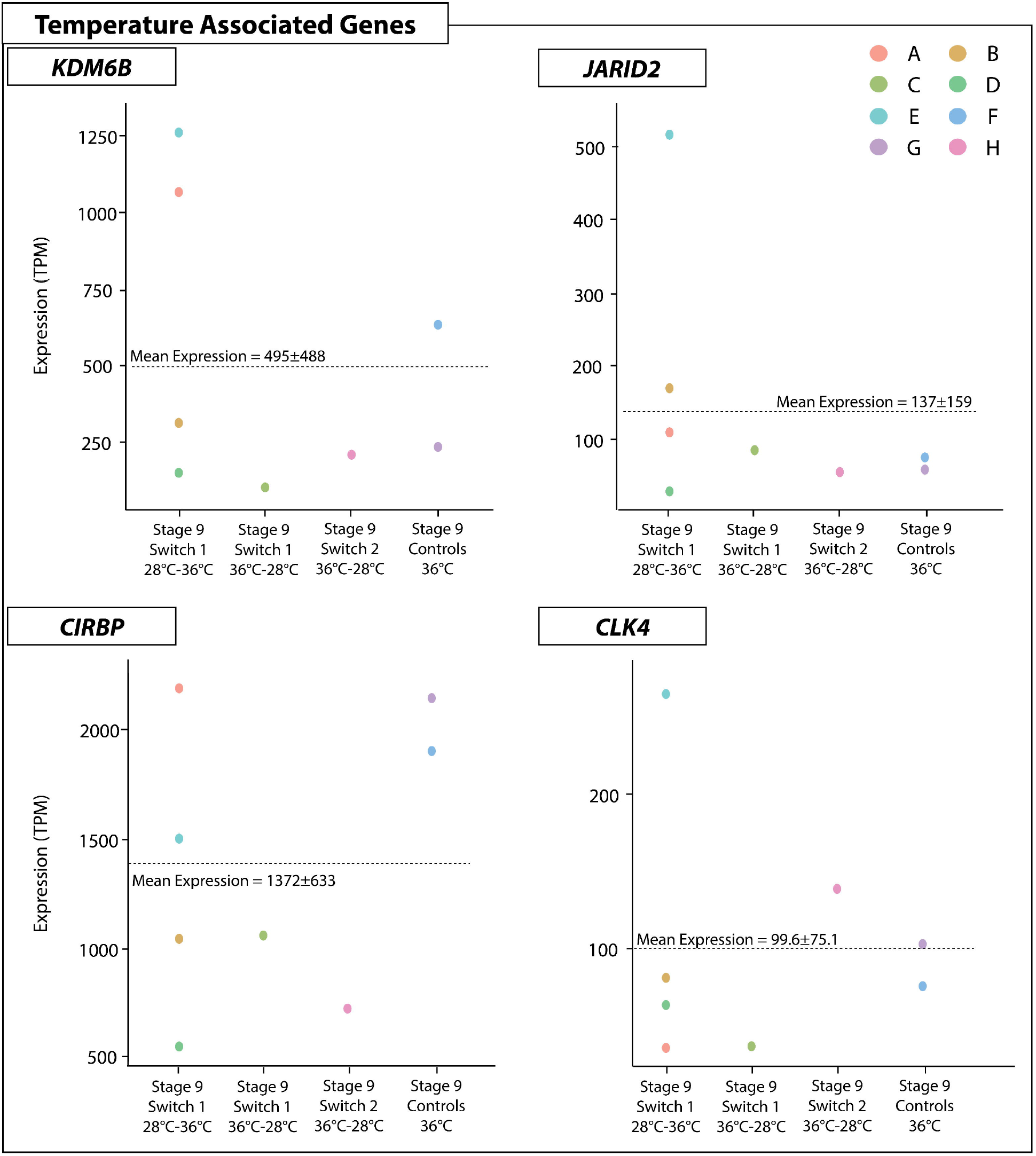
Expression (TPM, transcripts per million) of four temperature associated genes for all samples with ovotestes produced from different switching conditions (x-axis). Each point represents an individual (by colour) to depict the range of expression values different individuals can exhibit for a given gene. Sample IDs correspond to those provided in Supplementary File S13, and are as follows: A) 3603zz_18_1_4, B) 3603zz_18_1_7, C) 3603zz_18_1_8, D) 3603zz_18_2_4, E) 3632zz_18_2_9, F) 3632zz_18_2_19, G) 3632zz_18_2_20, H) 3232zz_18_1_19.

Inter-individual variation may be explained by differing responses to temperature switches relating to an individual’s sensitivity to the environment. This certainly is likely playing a role in the patterns observed in the expression of these genes, as not all individuals respond to the treatments in the same way. These results are also likely influenced by the inter-individual variation in the cellular makeup of ovotestes.

### Genes uniquely associated with ovotestes

In order to better understand the unique transcriptional profiles of ovotestes, differential gene expression analysis was conducted between ovotestes produced at stage 9 up-switched at stage 6 and the ovotestes from the 36°C controls, and control testes and ovaries both at stage 9. Ovaries from the control incubations were grouped with pre-switch ovaries at stage 9 from the mid developmental switch regime. The differentially expressed genes between ovotestes and testes, and ovotestes and ovaries can reveal the differences between the phenotypes, and provide clues to which phenotype the ovotestes more closely resemble, and where on the spectrum they lie. Then by taking the genes upregulated in ovotestes from both datasets it is possible to determine genes that are uniquely associated with ovotestes, and not with either testes or ovaries.

#### Differentiation gene expression between ovotestes, and ovaries and testes

There were many more genes differentially expressed between testes and ovotestes (not including testis like ovotestes) than there were between ovaries and ovotestes, suggesting that the expression profiles of ovotestes are typically more similar to ovaries than they are to testes. A total of 579 genes were upregulated in testes compared with ovotestes, and 557 in ovotestes compared with testes (Supplemental Table S8). Whereas 132 genes were upregulated in ovaries compared with ovotestes, and 97 were upregulated in ovotestes compared with ovaries (Supplemental Table S9).

A large number of genes were differentially expressed between ovotestes and testes. Various male specific genes were upregulated in testes compared with ovotestes, including *GHRH, AMH, DMRT1, HSD17B3, WIF1, SFRP2, WNT6, FZD6, FRZB, DLK2*, and *WLS*. In ovotestes, female related genes *CYP17A1* and *CYP19A1* were upregulated, as was *STAR, FZD4, PGR, DHRS3, SOX4, SOX12*, and *GATA2*. Two TRP channels were differentially expressed between testes and ovotestes. *TRPV4* was upregulated in testes, whereas *TRPV1* was upregulated in ovotestes. These two genes have high sequence similarity and are highly sensitive to temperature (34). A variety of stress response genes were differentially expressed between testes and ovotestes. Genes upregulated in testes included heat shock genes *HSPB2, DNAJB11, DNAJC15* and *DNAJB9*, antioxidant genes *TXNDC5, PRDX4*, and *SOD2*. In ovotestes, *CLK4*, a temperature sensitive splicing regulator, was upregulated alongside *STAT2* and *JAK3*, components of the JAK-STAT pathway, *RELA* and *IKBKE* which are part of the NF-kB pathway, as well as *JARID2* and *KDM6B*, chromatin remodelling genes associated with sex reversal and TSD. *CRH*, a gene involved in the hormonal stress response, and previously associated with sex reversal in adults (10), was also upregulated.

Far fewer genes were differentially expressed between ovaries and ovotestes than testes and ovotestes. Genes of note upregulated in ovaries cf ovotestes included antioxidant gene *TXNDC5*, which was also upregulated in testes cf ovotestes, oxidative stress gene *OSGIN1*, and splicing regulator *NOVA1*. In ovotestes, another member of the same lysine demethylase family as *KDM6B, KDM5C*, was upregulated in comparison with ovaries, as was a different TRP channel, *TRPM3*. Oxidative stress genes *SQOR* and *GADD45G* were also upregulated cf ovaries. Unlike what was observed in the differential expression analysis between ovotestes and testes, no sex related genes were differentially expressed between ovaries and ovotestes. This implies that the transcriptional profiles of ovotestes are more similar to ovaries than to testes, despite the propensity for these gonads to differentiate as testes later in development following exposure to the same temperature regimes that produced these ovotestes.

#### Genes uniquely upregulated in ovotestes compared with ovaries and testes

Of the 689 genes upregulated in ovotestes across both datasets, 36 genes were uniquely upregulated in ovotestes when compared with both ovaries and testes (Supplemental Table S10). Within the 36 genes uniquely upregulated in ovotestes cf both testes and ovaries (Fig. 9), are two circadian rhythm genes, *PER1* and *CIART*. Both genes are circadian pacemakers in the mammalian brain, and are transcribed in a cyclical pattern in the suprachiasmatic nucleus (35–37). Other organs in the body have peripheral circadian clocks, including mammalian ovaries, where *PER1* expression cycles during a 24h period in the steroidogenic cells (38). However, mammalian testes do not appear to possess a peripheral clock, and *PER1* expression appears to be developmental only (38,39). *CIART* has no described role in the gonads. It has previously been shown to be upregulated in the brains adult sex reversed female *P. vitticeps* (10). An experiment assessing thermal adaptation in three Anole species found that both *PER1* and *CIART* were upregulated in brain in response to temperature (40). The role these genes may play, or the influence circadian oscillations may have more broadly, in *P. vitticeps* ovotestes is unknown and requires further investigation.

**Figure 9:**
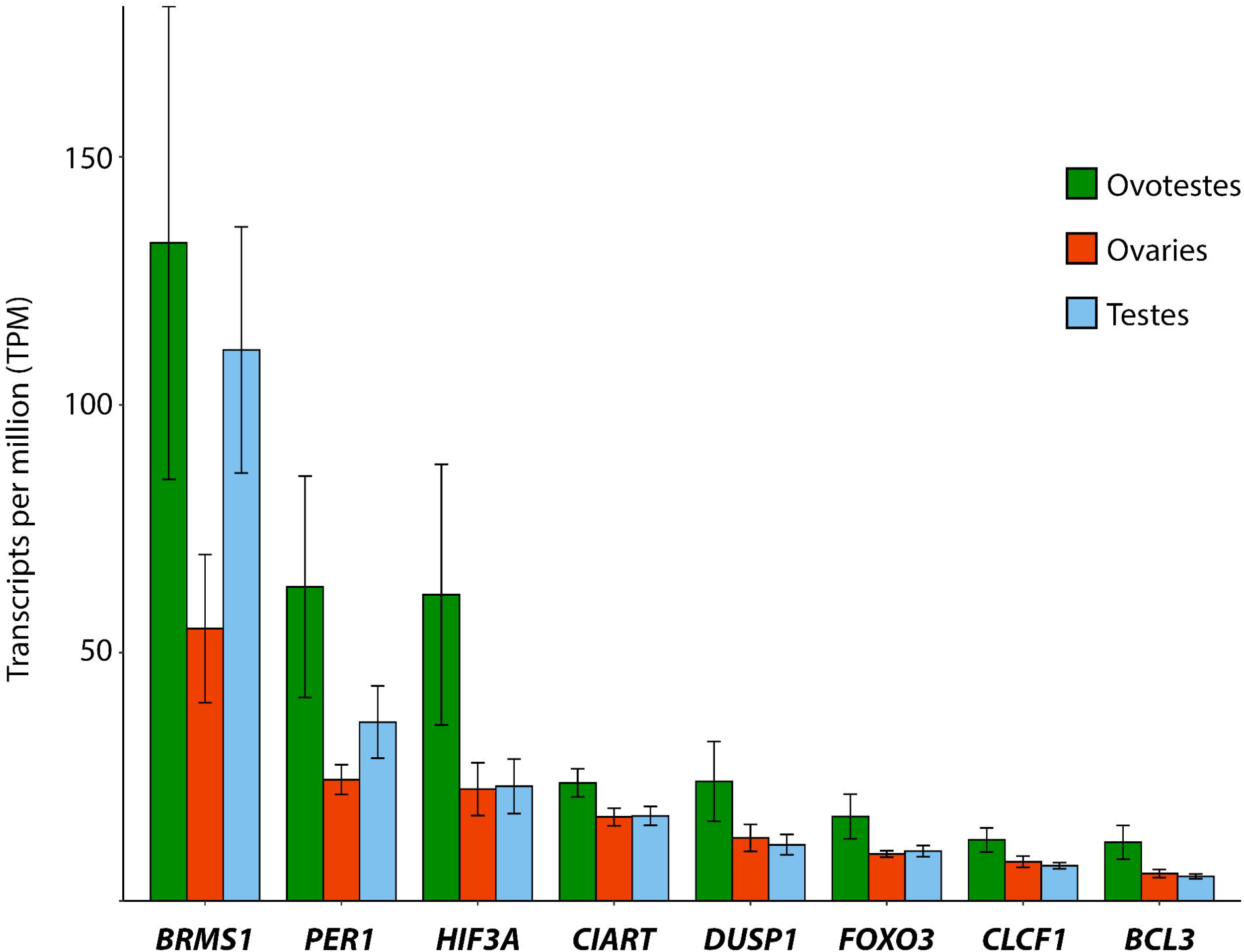
Expression (transcripts per million, TPM) for a subset of genes uniquely upregulated in ovotestes mentioned in the text. For the full list of genes see supplementary file S10.

Various genes associated with environmental stress pathways were uniquely upregulated in ovotestes. These included NF-kB pathway regulators *BCL3* and *BRMS1*, JAK-STAT pathway activator *CLCF1*, and *DUPS1* which regulates a MAP/ERK pathway. Also upregulated were the environmental stress genes *FOXO3* and *HIF3A*. Upregulation of these types of genes is to be expected at high incubation temperatures, however they were not differentially expressed in ovaries from the same temperature, suggesting they play a unique role in ovotestes.

## Discussion

Taken together, these findings reveal the complex effects of temperature on both gonadal morphology and gene expression in *P. vitticeps* during embryonic development. This study presents a novel approach that provides data revealing new insights into temperature responses in *P. vitticeps*, but that also has implications for vertebrates more broadly. By combining gonad histology with matched transcriptomes, this is the first study to our knowledge to sequence reptile ovotestes, and to be able to analyse gene expression profiles with validated morphology. These findings also provide new evidence supporting the role of CaRe mechanisms in sensing and transducing temperature cues in sex determination cascades.

The gonadal phenotypes produced by temperature switches at different developmental stages revealed that the gonads are responsive to temperature at stage 6, but are insensitive to temperature shifts by stage 9. The results from the early developmental switches suggest that a thermal signal of considerable strength is required to override male development and cause sex reversal. The testes-like phenotypes observed in the stage 6 samples at 36°C also suggests a propensity for male-ness even in the absence of exposure to 28°C. These results suggest, as has previously been hypothesised, that male development can be considered as the developmental default for *P. vitticeps* (21). Thus understanding sex reversal as a process by which the male pathway must be actively repressed or overridden by the female pathway can help explain why even short exposure to the male producing temperature can be sufficient to cause testes differentiation. This may also explain instances where sex reversal does not occur at high temperatures (“resistance” to reversal). In these individuals there may be differences in temperature sensitivity and/or the threshold of expression required to initiate sex reversal, be that repression of male genes, activation of female genes, or both (Fig. 10).

**Figure 10:**
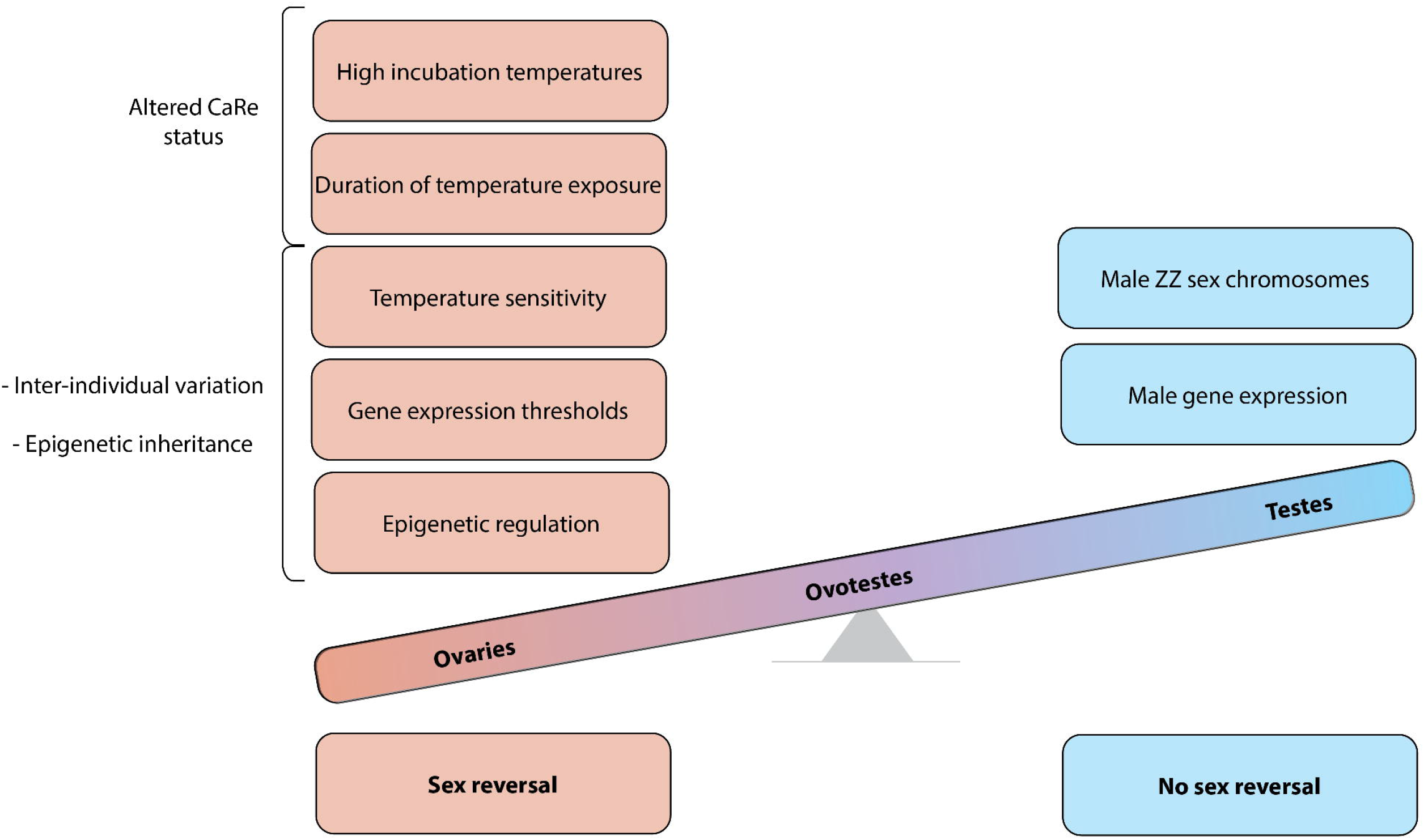
Schematic representation of the “threshold model” of sex reversal illustrating the many factors involved in initiating and maintaining sex reversal compared with male development. While male development readily progresses due to the presence of male sex chromosomes and the expression of male specific genes, the initiation of sex reversal relies on the compounding influence of many factors. Ovotestes are produced as an intermediate phenotype due to the competing influences of male and female sex determination pathways.

The transcriptomes of ovotestes show that like what is observed morphologically, ovotestes do indeed exhibit a mixed expression profile of both male and female specific genes. We further showed that there is considerable inter-individual variation in the expression profiles of ovotestes, such that they exist along a spectrum of male-ness to female-ness. This is also reflected in the phenotypes; some individuals may possess more obviously differentiated seminiferous tubules than others, or a thicker cortex with less well formed tubules. The transcriptional profile of ovotestes was found to have 32 unique genes with a variety of functions. Most are not normally associated with gonadal development or function, so require further study to understand the role the may play in sex reversal. None of these 32 genes have previously been associated with TSD in reptiles, or other ESD systems in different vertebrate groups. Due to a lack of research in this area, it is currently unclear whether this is because these genes are unique to the ovotestes phenotype, or if they may be more commonly associated with ESD systems in different species that have not yet been studied. This provides an important foundation for future research, as new species with sex reversal will likely be uncovered (41).

Analysis of gene expression changes following temperature switches showed in many cases there was expression of genes associated with the sex opposite to that seen morphologically in the same individual. This implies there may be delayed upregulation of genes from the opposite sex following a temperature switch to the opposite temperature, which is not reflected in the phenotype. This adds further weight to the hypothesis that various thresholds (temperature exposure, duration of exposure to that temperature, sensitivity to these changes, sufficiently high levels of female gene expression and/or sufficiently low levels of male gene expression) need to be surpassed for sex reversal to occur. The understanding provided by the CaRe model implies that an additional threshold may be the number of environmentally sensitive pathways that are activated by high incubation temperatures (5).

A threshold based understanding can explain some of the observed characteristics of sex reversal. This includes how rates of sex reversal increase as temperature increases: the more extreme the temperature, the more likely it is for everything to be pushed above these thresholds. Epigenetic inheritance, or genetic variants influencing these thresholds can also explain how the offspring of sex reversed mothers reverse more readily that those produced from normal mothers. It is likely that heritable epigenetic elements of sex reversal causes increased sensitivity to temperature and/or a lowering of the various thresholds required for sex reversal.

This threshold hypothesis also implies that sex reversal is inherently difficult to produce, as a variety of molecular hurdles need to be cleared in order for sex reversal to be initiated and maintained (Fig. 10). If this is indeed the case, why then does sex reversal occur at all? Current understanding of ESD systems, and in particular TSD, states that there must be fitness differences between the sexes under different environmental conditions to drive the evolution of these systems. In the case of *P. vitticeps*, additional complexity is introduced by the presence of sex chromosomes, as well as interaction with incubation temperature via epigenetic mechanisms and/or genetic variants influencing sex determination. Studies on wild populations of *P. vitticeps* have shown that sex reversal is not widespread, occurring in specific areas which do not experience the hottest temperatures (4). As has been hypothesised, this implies that local adaptation can occur, limiting the rates of sex reversal, perhaps conferring a selective benefit by ensuring the W chromosome is maintained in the population (42).

The data provided by our study has given new understanding to the influences of temperature on sex determination and differentiation in *P. vitticeps*, and also lays an important foundation upon which future work can be built. The unique combination of temperature switching with gonad morphology and gene expression analysis, has provided new insights into not only the complex effects of temperature on sex determination and differentiation in *P. vitticeps*. The results from this study also have implications for understanding environmentally sensitive sex determination systems in other vertebrate species more broadly. Ultimately, the more research that is conducted into ESD systems, more questions and curiosities are uncovered, but we continue to move closer to a more complete understanding of the interactions between sex and temperature in environmentally sensitive species.

## Materials and Methods

### Egg incubation and experimental design

Eggs were obtained from the University of Canberra breeding colony during the 2018-19 breeding season in accordance with approved animal ethics procedures (Project 270). Breeding groups comprised of three sex reversed ZZf females to one male, providing some control over the effects of male genotype. Sex reversal was validated using standard genotyping procedures (2), and phenotype and reproductive output. Females were allowed to lay naturally, and eggs were collected at lay or within two hours of lay. Following inspection for egg viability (presence of vasculature inside the eggshell), eggs were incubated in temperature-controlled incubator (±1°C) on damp vermiculite (four parts water to five parts water by weight).

Eggs from clutches of ZZf mothers were randomly split between 36°C and 28°C incubation temperatures, and then assigned randomly to one of four experimental regimes. For the offspring of sex reversed ZZf females, 36°C produces 100% females, and 28°C produces 100% males (2). For Switch 1, a subset of eggs were sampled at stage 6 from both incubation temperatures, and the remaining eggs were switched to the opposite incubation temperature, and sampled post-switch at stage 9 and 12. For Switch 2, a subset of eggs were sampled at stage 9 from both incubation temperatures, the remaining eggs switched to the opposite incubation temperature, and sampled post-switch at stage 12 and 15. For Switch 3, a subset of eggs were sampled at stage 12 from both incubation temperatures, the remaining eggs were switched to the opposite incubation temperature, and sampled post-switch at stage 15. Additional controls were incubated at constant temperatures (either 36°C or 28°C) and sampled at each of stage 9, 12, and 15 (Fig. 1). All embryos were staged according to the system developed for *P. vitticeps* in order to target particular developmental stages of interest (21,24). These developmental stages were selected based on previous research characterising gonadal development during sex reversal in *P. vitticeps* (21). At stage 6 the gonads are bipotential, and stage 9 is a period of developmental antagonism frequently characterised by the presence of ovotestes. Gonad differentiation occurs during stage 12, and by stage 15 the gonads have completely differentiated. Final sample sizes are given in Fig. 1 and Supplemental Table S13.

Embryos were sampled at the targeted developmental stages, taking into account differing developmental rates at the two incubation temperatures. Embryos were euthanised via intracranial injection of sodium pentobarbitone (60mg/ml in isotonic saline). For each embryo, one whole gonad was dissected from the surrounding mesonephros and snap frozen in liquid nitrogen for sequencing. The other whole gonad and mesonephros was dissected and preserved in neutral buffered formalin for histological processing. This design matches the phenotypic and genotypic data for each individual. This was also used to validate which samples possessed ovotestes prior to sequencing at stage 9, as ovotestes frequency is not 100%.

### Histology

Samples were processed for histology following procedures previously used in *P. vitticeps* described in (21). Briefly, samples were preserved in neutral buffered formalin before being transferred to 70% ethanol to improve tissue stability. All tissue processing was conducted at the University of Queensland’s School of Biomedical Sciences Histology facility. Samples were dehydrated, embedded in wax, and sectioned 6µm thick. Slides were stained using hematoxylin and eosin. The slides were scanned at 20x using the Aperio slide scanning system, and visualised using the Leica ImageScope program. Gonadal phenotypes were characterised using standard characteristics previously used for the species, and that are common to vertebrate gonads (21). For samples collected at stage 9, ovotestes phenotype was confirmed prior to sequencing.

### RNA extraction and sequencing

RNA from the isolated gonadal tissues was extracted in randomised batches using the Qiagen RNeasy Micro Kit (cat. No. 74004) according to manufacturer protocols. RNA was eluted in 14µl of RNAase free water and frozen at -80°C prior to sequencing. All samples were prepared for sequencing at the Australian Genome Research Facility (Melbourne, Australia). Sequencing libraries were prepared in randomised batch, and sequenced using the Novaseq 6000 platform (Illumina).

### Expression profiling and differential gene expression analysis

Paired-end RNA-seq libraries (.fastq format) were prepared using the sample pipeline that was established previously for *P. vitticeps* (8). Briefly, libraries were trimmed using trim_galore with default setting, and then aligned with the *P. vitticeps* reference NCBI genome using STAR. PCR duplicates and non-unique alignments were removed using samtools. Gene expression counts and normalised gene expression (transcripts per million, TPM) were determined using RSEM. Raw count and TPM files are provided (Supplemental Tables S11 and S12 respectively). Sample metadata is provided in Supplemental Table S13.

Differential gene expression analysis was conducted on raw count data using EdgeR. One sample identified as an outlier following PCA analysis was removed from the dataset (Fig. 3). Genes with fewer than 10 counts across three samples were removed as lowly expressed. Samples were grouped according to their position in the experimental design (samples from the first switch regime at 36°C sampled at stage 6 was the first group, and so on. See Fig. 1). Samples were also grouped according to temperature and phenotype. Full details of the EdgeR parameters used are described in (8). A p-value cut-off of ≤0.01 and a log_2_ fold-change threshold of 1 or -1 was applied to all contrasts. To determine genes uniquely expressed in ovotestes, samples from different groups at stage 9 were combined to increase sample sizes. The ovotestes samples included those produced from the control incubations at 36°C and from the switch 1 regime (switched from 28°C to 36°C at stage 6, sampled at stage 9). Stage 9 ovaries were combined from the control incubations, and from the pre-switch stage 9 sampling at 36°C in the second switch regime. Stage 9 testes were combined between the control 28°C and pre-switch stage 9 samples from the second switch regime.

## Acknowledgements

We thank Jin Dai for his assistance with RNA extractions, and Dr Ira Deveson for his assistance with processing the raw sequencing data. We thank Dr Wendy Ruscoe and Jacqui Richardson for their animal husbandry expertise.

## Author Contributions

S.L.W conducted all experimental procedures, data analysis, and led the preparation of the manuscript. A.G and S.L.W designed the experiment with assistance from C.E.H. A.G and C.E.H contributed to writing and editing the manuscript.

## Supplementary Materials

Supplemental Table S1: Genes differentially expressed between testes-like gonads at 36°C and testes at 28°C from the early developmental switch regime sampled at stage 6. A log-fold change cut-off of -1 to 1 and P-value cut-off of 0.01 was applied. The “direction” column indicates genes that were upregulated in the testes-like gonads compared with testes (negative log-fold changes), and those that were upregulated in the testes compared with testes-like gonads (positive log-fold changes).

Supplemental Table S2: Genes differentially expressed between the testes at 36°C and testes at 28°C from the early developmental switch regime sampled at stage 12. A log-fold change cut-off of -1 to 1 and P-value cut-off of 0.01 was applied. The “direction” column indicates genes that were upregulated in the 28°C testes compared with 36°C testes (negative log-fold changes), and those that were upregulated in the 36°C testes compared with 28°C testes (positive log-fold changes).

Supplemental Table S3: Genes differentially expressed between the testes at 28°C and ovaries at 36°C from the mid developmental switch regime sampled at stage 9. A log-fold change cut-off of -1 to 1 and P-value cut-off of 0.01 was applied. The “direction” column indicates genes that were upregulated in the 28°C testes compared with 36°C ovaries (negative log-fold changes), and those that were upregulated in the 36°C ovaries compared with 28°C testes (positive log-fold changes).

Supplemental Table S4: Genes differentially expressed between ovotestes at 28°C and ovotestes at 36°C from the early developmental switch regime sampled at stage 9. A log-fold change cut-off of -1 to 1 and P-value cut-off of 0.01 was applied. The “direction” column indicates genes that were upregulated in the 36°C ovotestes compared with 28°C ovotestes (negative log-fold changes), and those that were upregulated in the 28°C ovotestes compared with 36°C ovotestes (positive log-fold changes).

Supplemental Table S5: Genes differentially expressed between control ovotestes at 36°C and ovotestes from the early developmental switch regime up-switched from 28°C to 36°C sampled at stage 9. A log-fold change cut-off of -1 to 1 and P-value cut-off of 0.01 was applied. The “direction” column indicates genes that were upregulated in the up-switched ovotestes compared with control ovotestes (negative log-fold changes), and those that were upregulated in the control ovotestes compared with up-switched ovotestes (positive log-fold changes).

Supplemental Table S6: Genes differentially expressed between stage 9 ovotestes and stage 12 testes at 36°C from the early developmental switch regime. A log-fold change cut-off of -1 to 1 and P-value cut-off of 0.01 was applied. The “direction” column indicates genes that were upregulated in the testes compared with ovotestes (negative log-fold changes), and those that were upregulated in ovotestes compared with testes (positive log-fold changes).

Supplemental Table S7: Genes differentially expressed between stage 9 up-switched ovotestes (early developmental switch regime) and stage 9 pre-switch ovaries at 36°C (mid developmental switch regime). A log-fold change cut-off of -1 to 1 and P-value cut-off of 0.01 was applied. The “direction” column indicates genes that were upregulated in the ovaries compared with ovotestes (negative log-fold changes), and those that were upregulated in ovotestes compared with ovaries (positive log-fold changes).

Supplemental Table S8: Genes differentially expressed between stage 9 ovotestes and stage 9 control testes. Ovotestes produced at stage 9 up-switched at stage 6 and the ovotestes from the 36°C controls were grouped. A log-fold change cut-off of -1 to 1 and P-value cut-off of 0.01 was applied. The “direction” column indicates genes that were upregulated in the testes compared with ovotestes (negative log-fold changes), and those that were upregulated in ovotestes compared with testes (positive log-fold changes).

Supplemental Table S9: Genes differentially expressed between stage 9 ovotestes and stage 9 ovaries (controls and pre-switch stage 9 36°C from the mid-development switch regime samples were grouped). Ovotestes produced at stage 9 up-switched at stage 6 and the ovotestes from the 36°C controls were grouped. A log-fold change cut-off of -1 to 1 and P-value cut-off of 0.01 was applied. The “direction” column indicates genes that were upregulated in the ovaries compared with ovotestes (negative log-fold changes), and those that were upregulated in ovotestes compared with ovaries (positive log-fold changes).

Supplemental Table S10: Genes uniquely upregulated in stage 9 ovotestes compared with both ovaries and testes (also at stage 9). Ovotestes produced at stage 9 up-switched at stage 6 and the ovotestes from the 36°C controls were grouped. Stage 9 ovaries (controls and pre-switch stage 9 36°C from the mid-development switch regime) were grouped. Genes that were differentially between both comparisons (ovotestes vs ovaries, and ovotestes vs testes) were determined to be uniquely upregulated in ovotestes. A log-fold change cut-off of -1 to 1 and P-value cut-off of 0.01 was applied.

Supplemental Table S11: Raw counts for all samples (n = 58) for all genes (n = 19,284) prior to any filtering or outlier sample removal. Sample metadata is provided in Supplemental Table S13.

Supplemental Table S12: Raw expression values (TPM, transcripts per million) for all samples (n = 58) for all genes (n = 19,284) prior to any filtering or outlier sample removal. Sample metadata is provided in Supplemental Table S13.

Supplemental Table S13: Metadata for all samples in this study. The original sample ID is matched to the Data ID, which appears in the raw count (Supplemental Table S11) and raw expression (Supplemental Table S12) files. The parent IDs are provided, and the switch regime that each sample was used in is noted. The group column corresponds to the data ID, and where these groups are in the experiment design is provided in the “groups” sheet. This re-naming is to facilitate data analysis. The gonadal phenotype, and how the phenotype was validated (either via histology or expression profiles) is also provided.

